# Functional Equivalence of Gamma- and X-Ray Irradiation for Long-Term Hematopoietic and AML Transplant Outcomes in Mice

**DOI:** 10.64898/2025.12.15.694472

**Authors:** Shawn David, Griffin J. Nye, Ewelina Bolcun-Filas, Jennifer J. Trowbridge, Kira A. Young

**Author notes:** Co-Corresponding Authors: Jennifer J. Trowbridge and Kira A. Young, The Jackson Laboratory, 600 Main Street, Bar Harbor, Maine, USA 04609. **Competing Interests:** All authors declare no competing financial interests. **Author Contributions:** S.D. analyzed data and wrote the manuscript. G.J.N. generated data and edited the manuscript. E.B-F. obtained funding for the project and edited the manuscript. J.J.T. designed the study, analyzed data, and edited the manuscript. K.A.Y. designed the study, generated data, analyzed data, and edited the manuscript.

## Abstract

Total-body irradiation (TBI) is routinely used for myeloablation prior to mouse hematopoietic cell transplant. Widespread transition from ^137^Cs *γ*-irradiators to X-ray systems has raised questions about whether these modalities yield equivalent biological outcomes. Although prior studies compared *γ* and X-ray irradiation in healthy syngeneic transplants, their performance in reciprocal congenic models and in primary acute myeloid leukemia (AML) transplant remains unclear. Here, we systematically evaluated *γ* and X-ray irradiation across dose and dose-rate conditions, and tested dose equivalents in CD45.1/CD45.2 reciprocal transplants and in AML transplant models. While each modality exhibited distinct early effects, both ultimately supported comparable long-term donor chimerism in congenic transplants and equivalent AML engraftment, leukemic burden, and disease progression. These findings indicate that, with proper dose calibration, X-ray irradiation is a functionally effective alternative to *γ*-irradiation for normal and malignant transplant studies.

## Introduction

Ionizing radiation is considered the gold standard to ablate endogenous hematopoiesis for transplantation[1]. Increasing regulatory scrutiny, cost, and security burden associated with radioactive sources have driven many institutions to replace these with self-contained X-ray irradiators[1, 2]. Relatively few studies have systematically compared these two modalities in experimental transplant settings. One of the first direct comparisons reported that both sources can sufficiently ablate endogenous bone marrow, however, there were differences in lineage reconstitution[1]. B cell reconstitution was more robust following *γ*-irradiation, while X-ray-irradiated mice exhibited relatively higher T-cell reconstitution, albeit at the expense of decreased survival. More recently, comparison of gamma and X-ray irradiation revisited this issue using modern 350keV X-ray irradiation versus ^137^Cs *γ*-rays with dose titration. In this study, both modalities achieved near-complete donor chimerism, but only when doses were appropriately adjusted (e.g., 13Gy gamma versus ∼11Gy X-ray)[3]. These data support the feasibility of X-ray platforms but highlight that optimization is needed for experimental setup.

Here, we evaluated and optimized X-ray irradiation in two experimental contexts not previously reported in the literature. First, in reciprocal congenic transplants between C57BL/6J (CD45.2) and B6.SJL (CD45.1) which are frequently and interchangeably used in most mouse transplant studies, and second, in mouse primary AML cell transplant using *Dnmt3a;Npm1*-AML[4]. Such comparisons are highly relevant for the growing number of laboratories transitioning away from *γ*-irradiators toward X-ray platforms.

## Methods

### Animals

Female C57BL/6J (RRID:IMSR_JAX:00664) and B6.SJL-*Ptprc*^*a*^*Pepc*^*b*^/BoyJ (B6.SJL, RRID:IMSR_JAX:002014) mice were housed within The Jackson Laboratory (Bar Harbor, ME). All mice used in experiments were young adults (2-4 months old). Mice were maintained in a temperature controlled vivarium on a 12h:12h light dark cycle. All mouse work was approved by the Animal Care and Use Committee at The Jackson Laboratory.

### Total Body Irradiation

Bone marrow ablation was performed using a ^137^Cs *γ*-emitting irradiator with a 2x attenuator (Model Mark 1 Series) or an X-ray irradiator (XRad320, Precision X-Ray). Both irradiators were in the same room, at the same barrier level as the animals’ housing room. Mice received split doses (for lethal irradiation, 3-4 hours apart) or single doses (for sublethal irradiation) in mouse pie cages (MPC-1, Braintree Scientific) under constant rotation.

### Bone Marrow Transplant

Femurs, iliac crests and tibias were harvested from donor mice. Bones were crushed using a mortar and pestle and treated with 1x red blood cell (RBC) lysis buffer to isolate mononuclear cells (MNCs). Cryopreserved AML samples previously harvested from mice with monocytic *Dnmt3a*^R878H/+^ *Npm1*^cA/+^ AML [4] were recovered. Viable cell count was determined by trypan blue exclusion using a hemocytometer. 2x10^6^ viable MNCs or 500,000 viable AML cells were transplanted into each recipient mouse via retro-orbital injection under anesthesia. The health status of recipient mice was monitored daily and mice with body condition score ≤2.0 were euthanized.

### Peripheral Blood Analysis

Blood was collected via the retro-orbital sinus under anesthesia and RBCs were lysed using 1X RBC lysis buffer. MNCs were stained with fluorochrome-conjugated antibodies recognizing CD45.1 (clone A20), CD45.2 (clone 104), B220 (clone RA3-6B2), CD3e (clone 145-2C11), CD11b (clone M1/70), Ly6g (clone 1A8), Ly6c (clone HK1.4), F4/80 (clone BM8), and Ter119 (clone TER-119). DAPI was used as a viability stain. Data were captured by A5 Symphony SE (BD Biosciences) or LSRII (BD Biosciences) and analyzed on FlowJo V10.10.0 (FlowJo, LLC). Complete blood counts were performed using a XN-1000V (Sysmex).

### Data Analysis

Analyses were performed using Prism V10.4 (GraphPad).

## Results and Discussion

### 11Gy X-ray irradiation is functionally equivalent to 13Gy ^137^Cs irradiation in short-term hematopoietic reconstitution

We transplanted donor BM from C57BL/6J (CD45.2^+^) mice into age-matched congenic B6.SJL (CD45.1^+^) mice that had received split doses of 13Gy *γ*-irradiation from a ^137^Cs source, which is typically used as a lethal irradiation dose for BM transplant[3, 5], or split doses of 11Gy, 12Gy, 13Gy or 14Gy X-ray irradiation (Figure 1A). At 4 weeks post-transplant, mice in the 11Gy X-ray group had similar levels of donor-derived peripheral blood (PB) hematopoietic cells compared to the 13Gy *γ*-irradiated control group, whereas higher doses of X-ray irradiation (12Gy, 13Gy, 14Gy) resulted in higher levels of donor-derived PB (Figure 1B). Over the subsequent two weeks, all mice in the 13Gy and 14Gy X-ray groups developed severe ulcerative dermatitis and had to be sacrificed (Figure 1C). This is consistent with previously reported lethality of congenic BM transplant recipients receiving 13Gy X-ray irradiation[3]. At 8 weeks post-transplant, mice in the 11Gy X-ray group continued to have similar levels of donor-derived PB cells compared to 13Gy *γ* group, with 12Gy X-ray continuing to have higher levels (Figure 1D). Comparing multilineage engraftment at 8 weeks post-transplant, the 13Gy *γ*-irradiated, 11Gy X-ray and 12Gy X-ray groups had similar proportions of donor-derived myeloid, B and T cells (Figure 1E). Together, these data corroborate previous conclusions by an independent group that a similar degree of hematopoietic chimerism obtained with 13Gy *γ*-irradiation can be achieved by 11Gy X-ray irradiation[3].

**Figure 1.**
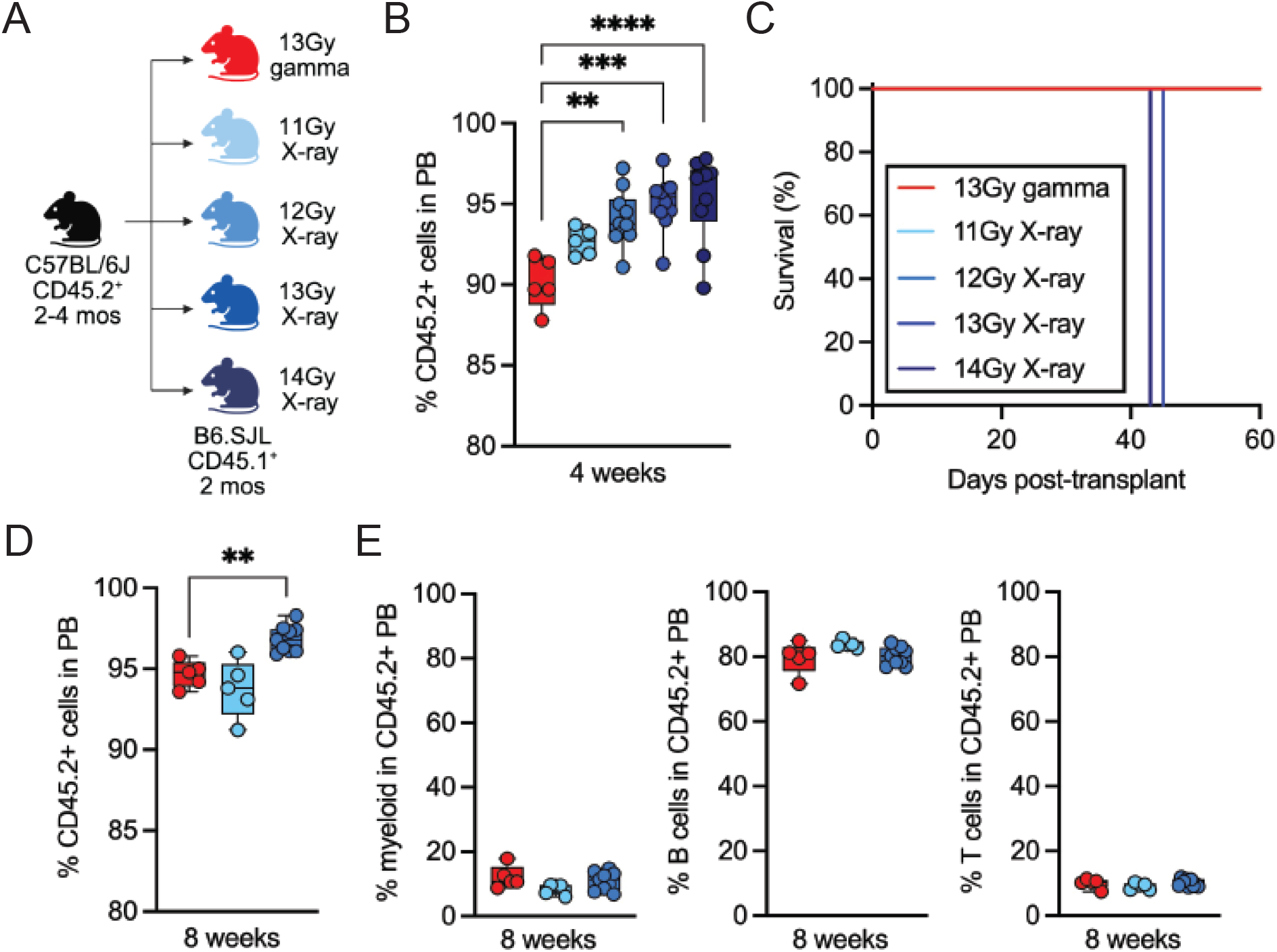
A lower X-ray irradiation dose versus gamma irradiation dose is needed to achieve similar levels of short-term hematopoietic chimerism. (**A**) Experimental design. (**B**) Frequency of donor-derived (CD45.2^+^) cells in the peripheral blood (PB) of recipient mice at 4 weeks post-transplant. (**C**) Survival of recipient mice over time. (**D**) Frequency of donor-derived cells in PB of recipient mice at 8 weeks post-transplant. (**E**) Lineage composition (myeloid, B cells, T cells) of donor-derived PB at 8 weeks post-transplant. (**B, D, E**) Dots represent individual recipient mice (*n* = 5-10 replicates per group). Data analyzed by one-way ANOVA with Dunnett’s multiple comparisons test. \*\**P*<0.01, \*\*\**P*<0.001, \*\*\*\**P*<0.0001.

Beyond total irradiation dose, an independent variable in BM transplant studies is dose rate. Dose rates have different impacts on distinct tissues [6] and previous BM transplant studies have shown that X-ray dose rate can impact hematopoietic chimerism[7, 8]. We tested dose rates of 0.72, 1.62 and 2.87 Gy/min to achieve the same total dose (11Gy) of X-ray irradiation. We observed no difference in proportion of donor-derived PB cells over 16 weeks post-transplant (Supplemental Figure 1A). Examining lineage composition, we observed a short-term increase in myeloid cell production at the expense of B cell production at the intermediate dose rate (1.62 Gy/min) that resolved in later time points (Supplemental Figure 1B-C). We also observed a delay in T cell production at the lowest dose rate (0.72 Gy/min) that resolved in later time points (Supplemental Figure 1D). These data show that dose rates may impact short-term lineage dynamics but do not significantly impact long-term multilineage hematopoiesis.

### 11Gy X-ray irradiation is functionally equivalent to 13Gy ^137^Cs irradiation in long-term hematopoietic reconstitution in the context of reciprocal congenic transplant

A common tool in experimental hematology is transplant of BM cells from mice on a C57BL/6J (CD45.2^+^) background into B6.SJL (CD45.1^+^) recipients, and mice on a B6.SJL (CD45.1^+^) background into C57BL/6J (CD45.2^+^) recipients, which maximizes flexibility in transplant design. Previous studies comparing *γ*-irradiation vs. X-ray irradiation used only the latter design (CD45.1^+^ into CD45.2^+^)[1, 3], which is less common when using genetically engineered mouse models as donors. Thus, we performed a direct comparison of both transplant designs. Donor C57BL/6J BM was transplanted into B6.SJL recipients, and donor B6.SJL BM was transplanted into C57BL/6J recipients, with both recipient groups receiving 11Gy X-ray irradiation (Figure 2A). As a control, donor B6.SJL BM was transplanted into 13Gy *γ*-irradiated C57BL/6J recipients. We observed differences in short-term hematopoiesis between these groups. At 4 weeks post-transplant, the 11Gy X-ray group (CD45.2^+^ into CD45.1^+^) had the highest donor-derived PB engraftment, followed by the 11Gy X-ray reciprocal group (CD45.1^+^ into CD45.2^+^), followed by the 13Gy *γ*-irradiated reciprocal group (CD45.1^+^ into CD45.2^+^) (Figure 2B). Despite this early difference, chimerism increased in all groups over time such that long-term hematopoiesis was not significantly different between groups. No differences were observed in reconstitution of the myeloid or B cell compartments between the groups (Figure 2C-D). With respect to T cell reconstitution, there was a delay in the 13Gy *γ*-irradiated group relative to both 11Gy X-ray groups (Figure 2E). This is consistent with a previous report that mice irradiated using X-ray have higher levels of T cell reconstitution compared to ^137^Cs[1]. These data show that performing reciprocal congenic transplants with X-ray irradiation may impact short-term hematopoietic reconstitution and lineage dynamics but does not significantly impact long-term multilineage hematopoiesis.

**Figure 2.**
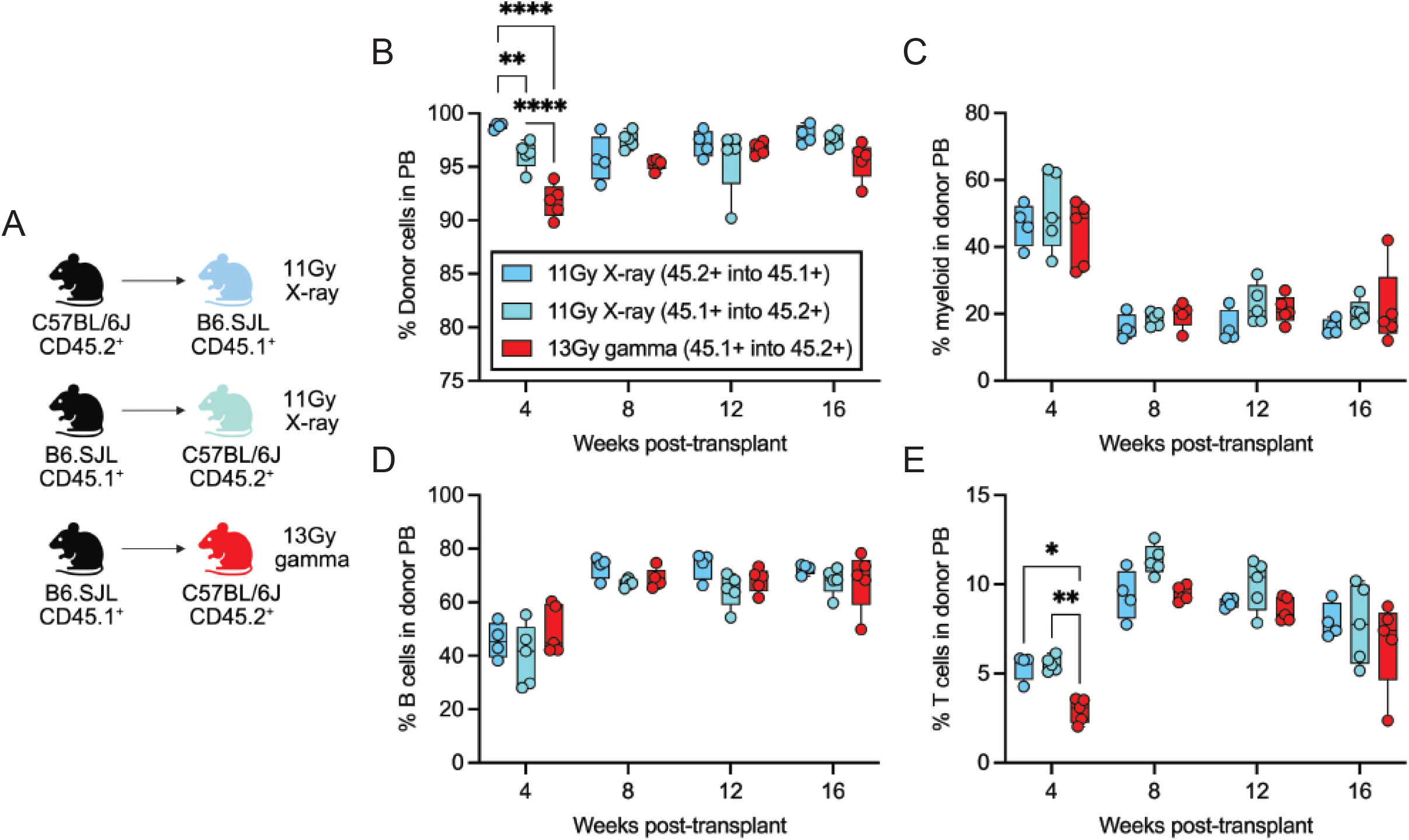
X-ray irradiation provides comparable long-term hematopoietic chimerism in standard reciprocal congenic transplant models. (**A**) Experimental design. (**B**) Frequency of donor-derived cells in the PB of recipient mice from 4 to 12 weeks post-transplant. (**C-E**) Proportion of myeloid (**C**), B cells (**D**) and T cells (**E**) in donor-derived PB from 4 to 12 weeks post-transplant. (**B-E**) Dots represent individual recipient mice (*n* = 4-5 replicates per group). Data analyzed by two-way ANOVA with Holm-Šídák’s multiple comparisons test.\**P*<0.05, \*\**P*<0.01, \*\*\*\**P*<0.0001.

### Sublethal doses of 5.5Gy X-ray and 6.5Gy ^137^Cs irradiation similarly condition mice for transplant and engraftment of primary mouse AML

Sublethal irradiation is commonly used in congenic transplant of primary mouse AML cells[4, 9, 10]. We compared X-ray versus *γ*-irradiation in this sublethal context using our previously established *Dnmt3a*^R878H/+^ *Npm1*^cA/+^ AML model[4]. Donor AML cells (CD45.2^+^) were transplanted into B6.SJL recipients receiving either 6.5Gy *γ*-irradiation or 5.5Gy X-ray irradiation (Figure 3A). At 4 weeks post-transplant, high donor myeloid cell chimerism was achieved in both conditioning regimens (Figure 4B-C). Both groups developed lethal AML in a similar timeframe (Figure 4D) and had comparable phenotypes including elevated white blood cell count and reduced red blood cell and platelet counts (Figure 4E). As expected, AML blasts in both groups were of the monocytic lineage (Figure 4F-G). These data show that sublethal X-ray irradiation results in comparable AML engraftment, leukemic burden, and disease progression.

**Figure 3.**
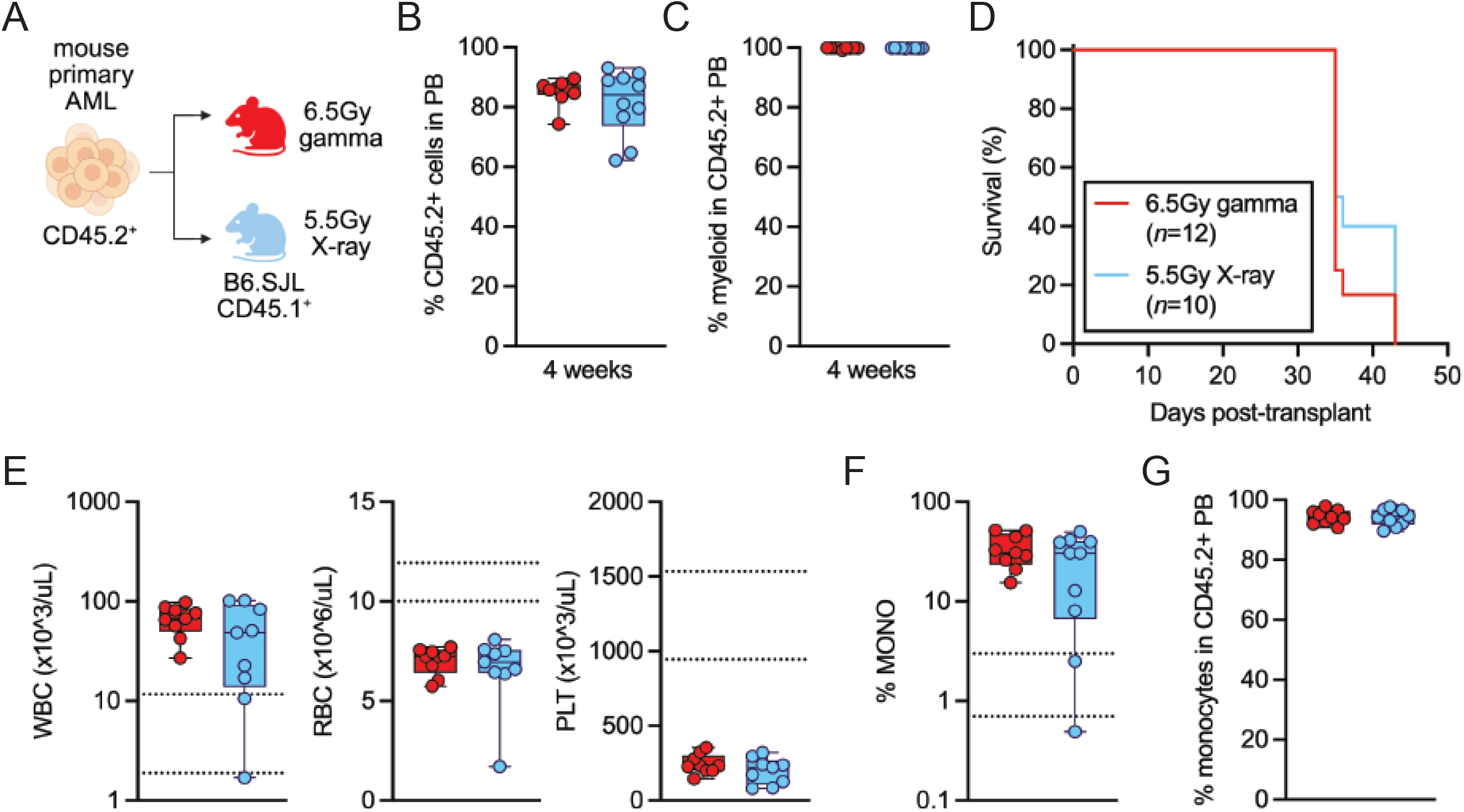
Sublethal gamma irradiation and X-ray irradiation efficiently condition recipient mice for transplant of primary AML cells. (**A**) Experimental design. (**B**) Frequency of donor-derived cells in the PB of recipient mice at 4 weeks post-transplant. (**C**) Proportion of myeloid cells in donor-derived PB at 4 weeks post-transplant. (**D**) Kaplan-Meyer survival curve of recipient mice. (**E**) White blood cell (WBC), red blood cell (RBC) and platelet (PLT) count in moribund recipient mice. (**F-G**) Frequency of monocytes in PB of moribund recipient mice using complete blood count (**F**) and flow cytometry (**G**) analysis. Dots represent individual recipient mice (*n* = 9-10 replicates per group). Data analyzed by unpaired, two-tailed *t* test and Log-rank (Mantel-Cox test).

## Acknowledgements

This work was supported by funding from the Pacific Northwest National Laboratory (PNNL) under Contract DE-AC05-76RL01830 with the U.S. Department of Energy. This study was funded in part by NIH grants R01DK118072, R01AG069010 and U01AG077925 to J.J.T. This work was supported in part by the NIH/NCI Cancer Center Support Grant P30CA034196. J.J.T. is a Scholar of Blood Cancer United. We thank Julie Alderman for X-ray irradiator support, and Jeremy Racine and Andy Greene for critical input into this project. We thank Mark K. Murphy and Maddison A. Heine (PNNL) for their assistance with dosimetry and the irradiator comparison studies. We thank the Scientific Services at The Jackson Laboratory including Shawnna Farley, Mark Warner, Jennifer Sargent and Muneer Hasham of the Xenografting and Live Imaging Service, Will Schott, Danielle Littlefield and Krystal Leigh-Brown of the Flow Cytometry Service, and Steven Ciciotte of the Clinical Chemistry Service. We thank all members of the Trowbridge Lab for project and manuscript feedback, and Nathan Boyer for experimental support.

## Figure Legends

**Supplemental Figure 1. Dose rate of X-ray irradiation does not impact long-term hematopoietic chimerism**. (**A**) Frequency of donor-derived cells in the PB of recipient mice from 4 to 16 weeks post-transplant. (**B-D**) Proportion of myeloid (**B**), B cells (**C**) and T cells (**D**) in donor-derived PB from 4 to 16 weeks post-transplant. Dots represent individual recipient mice (*n* = 5-8 replicates per group). Data analyzed by two-way ANOVA with Holm-Šídák’s multiple comparisons test. \**P*<0.05, \*\**P*<0.01.

