## Supplementary figures and images for "Functional Equivalence of Gamma- and X-Ray Irradiation for Long-Term Hematopoietic and AML Transplant Outcomes in Mice"

### Supplemental Figure 1

Supplemental Figure 1

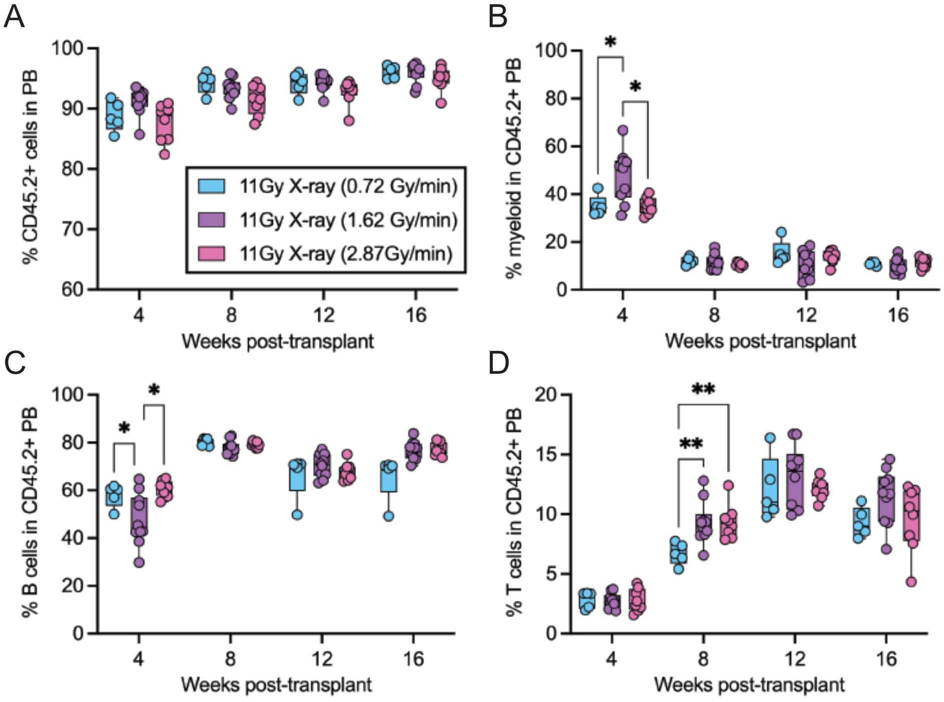
